## Supplementary Figures for "Chromosome-level genome assembly of *Torreya grandis* provides insights into the origin and evolution of gymnosperm-specific sciadonic acid biosynthesis"

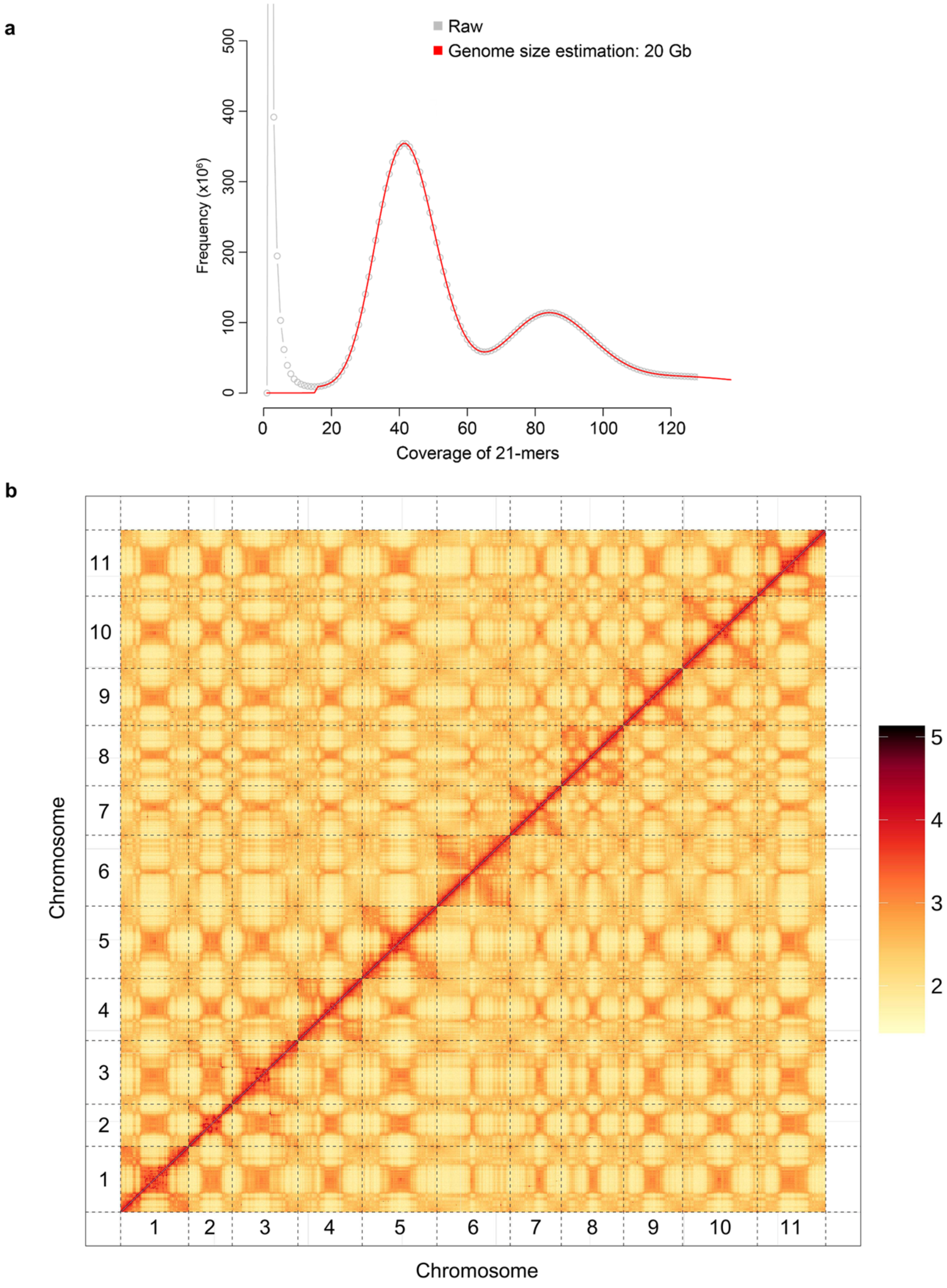


**Supplementary Figure 1. Size estimation and contig anchoring of the *T. grandis* genome.** (**a**) 21-mer spectrum of the *T. grandis* Illumina reads. Genome size was estimated using the FindGSE program (https://github.com/tiramisutes/findGSE). (**b**) Hi-C interaction map of *T. grandis* genome.

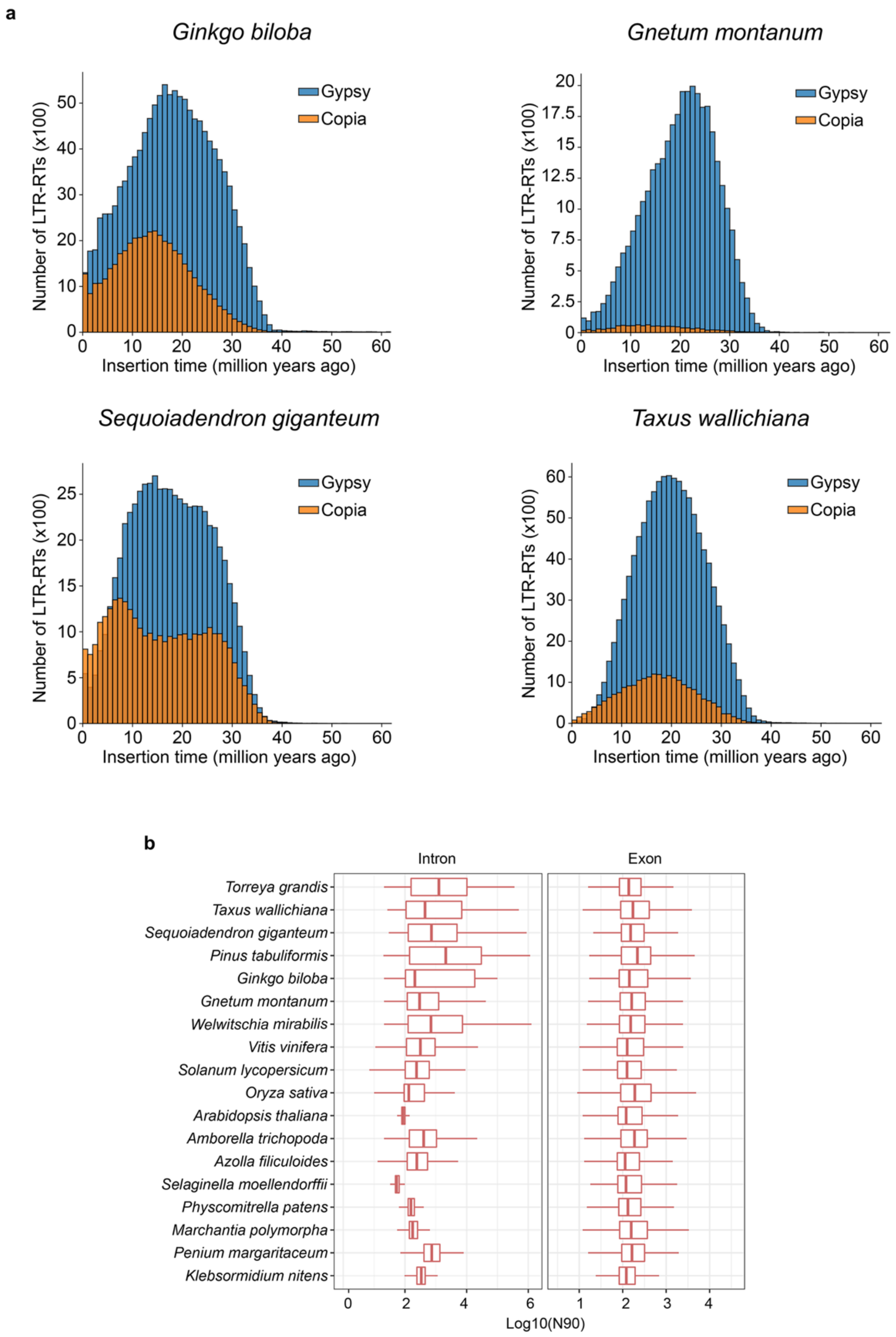


**Supplementary Figure 2. Distribution of LTR-RTs in selected genomes of gymnosperms**. (**a**) Expansion of *Copia* and *Gypsy* families in four selected genomes. (**b**) Boxplot of intron/exon lengths in different plant species. The N90 size of intron/exon for each gene was calculated and used for plot.

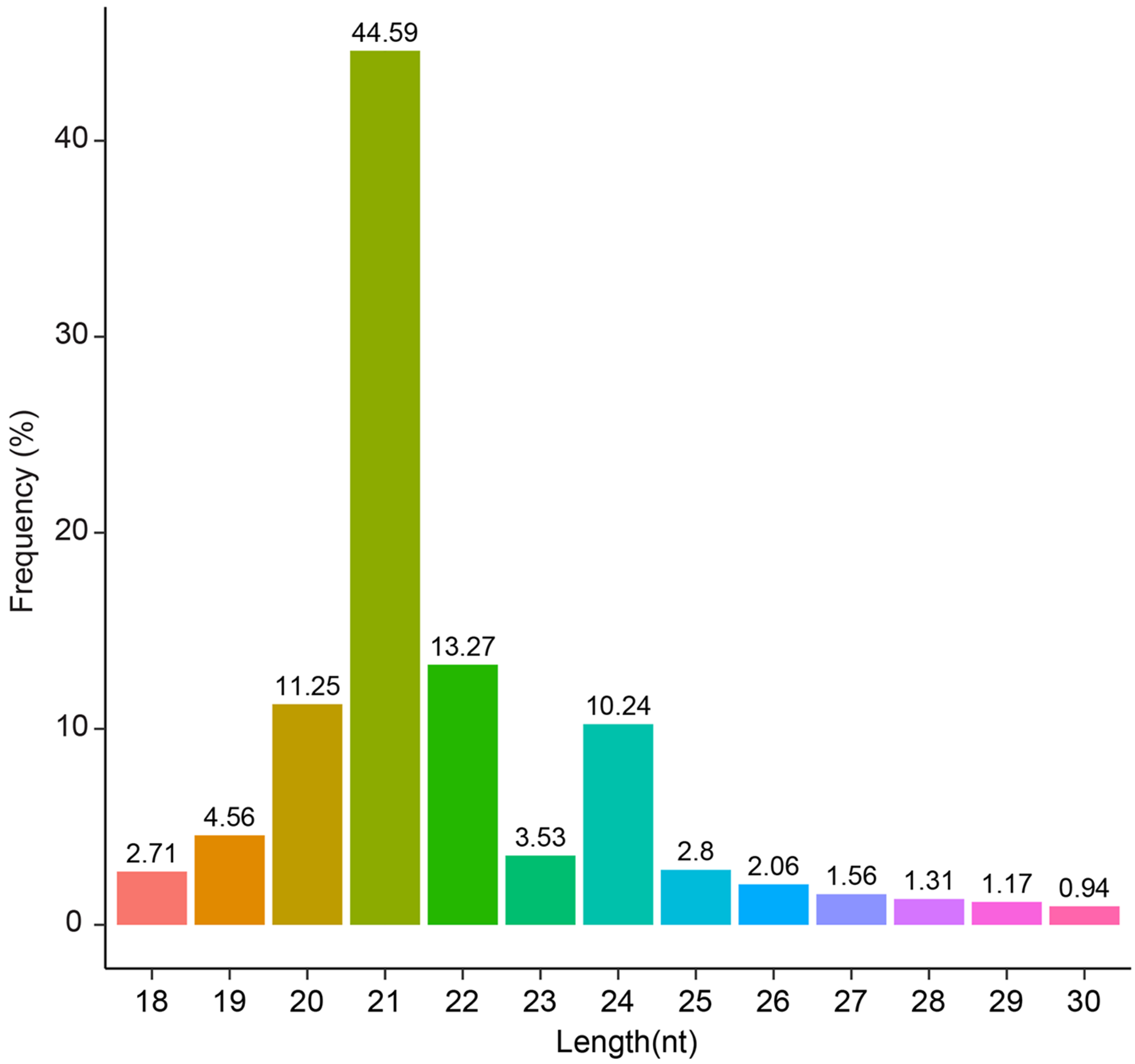


**Supplementary Figure 3. Size distribution of sRNAs in *T. grandis* leaves.** Percentages of sRNAs with different lengths are shown.

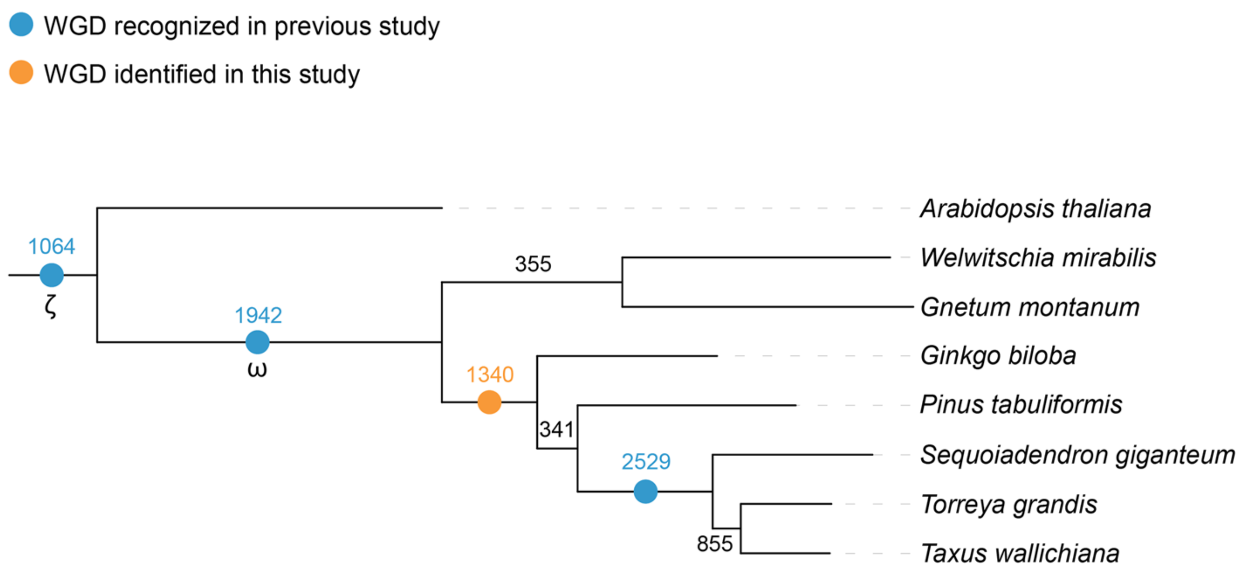


**Supplementary Figure 4. Tree based WGD inference.** Numbers on branches represent the quantity of duplicated gene families based on reconciliation of gene trees and species trees. Branches with high gene family duplication events usually imply WGDs.

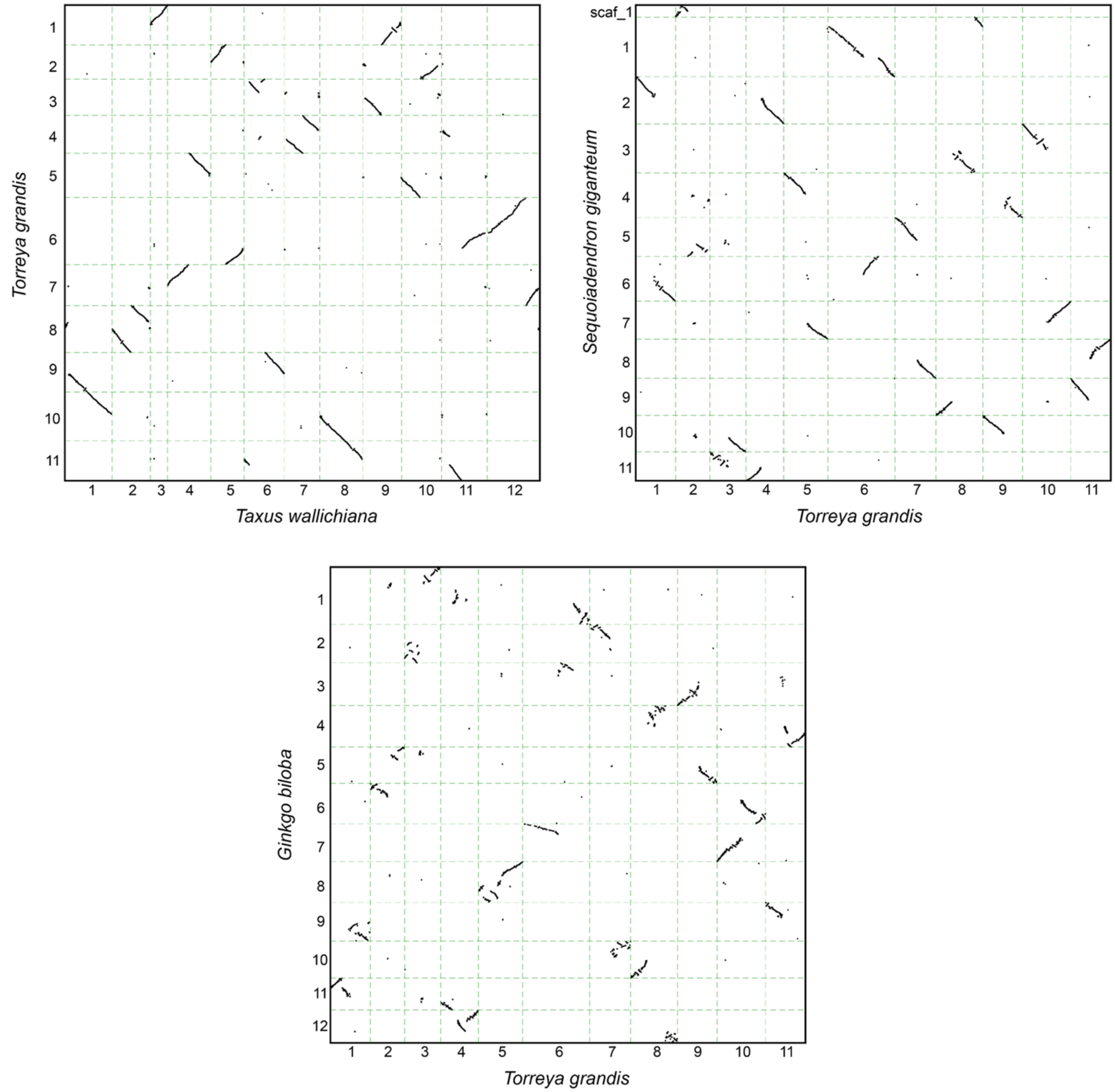


**Supplementary Figure 5. Collinearity between genomes of *T. grandis*, *S. gigantenum* and *G. biloba*.** Numbers on axis indicate chromosomes.

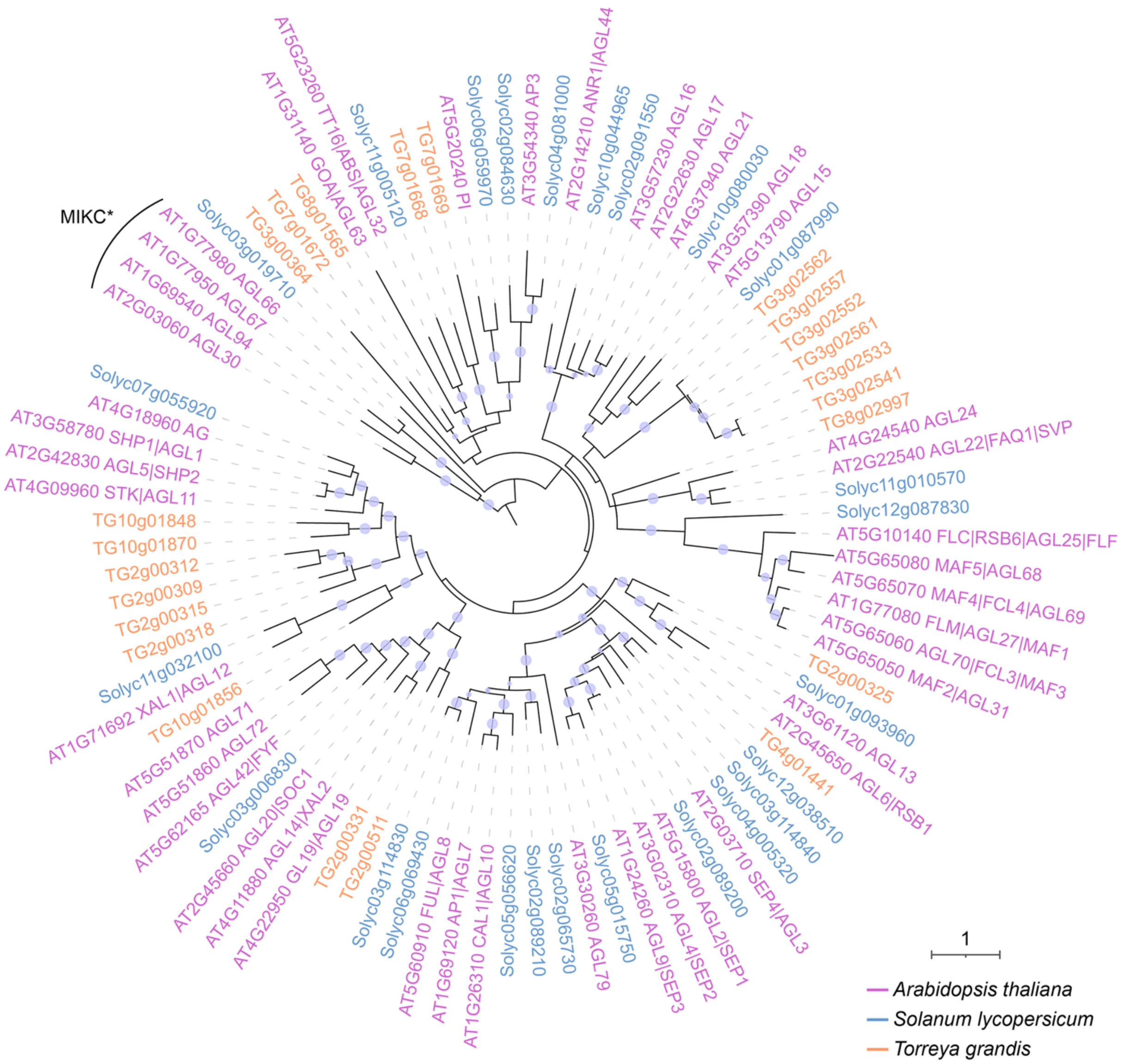


**Supplementary Figure 6. Phylogeny of MIKC$^{C}$ type MADS-box genes.** Protein sequences of MADS-box family genes were aligned using MAFFT with the linsi mode, and the phylogenetic tree was constructed using IQ-Tree with the best-fitting model (JTT+F+I+G4) and 1000 bootstrap replicates. Branches with bootstrap support greater than 80% were labeled with light blue pies. Tree was rooted with MIKC$^{*}$ type MADS-box genes.

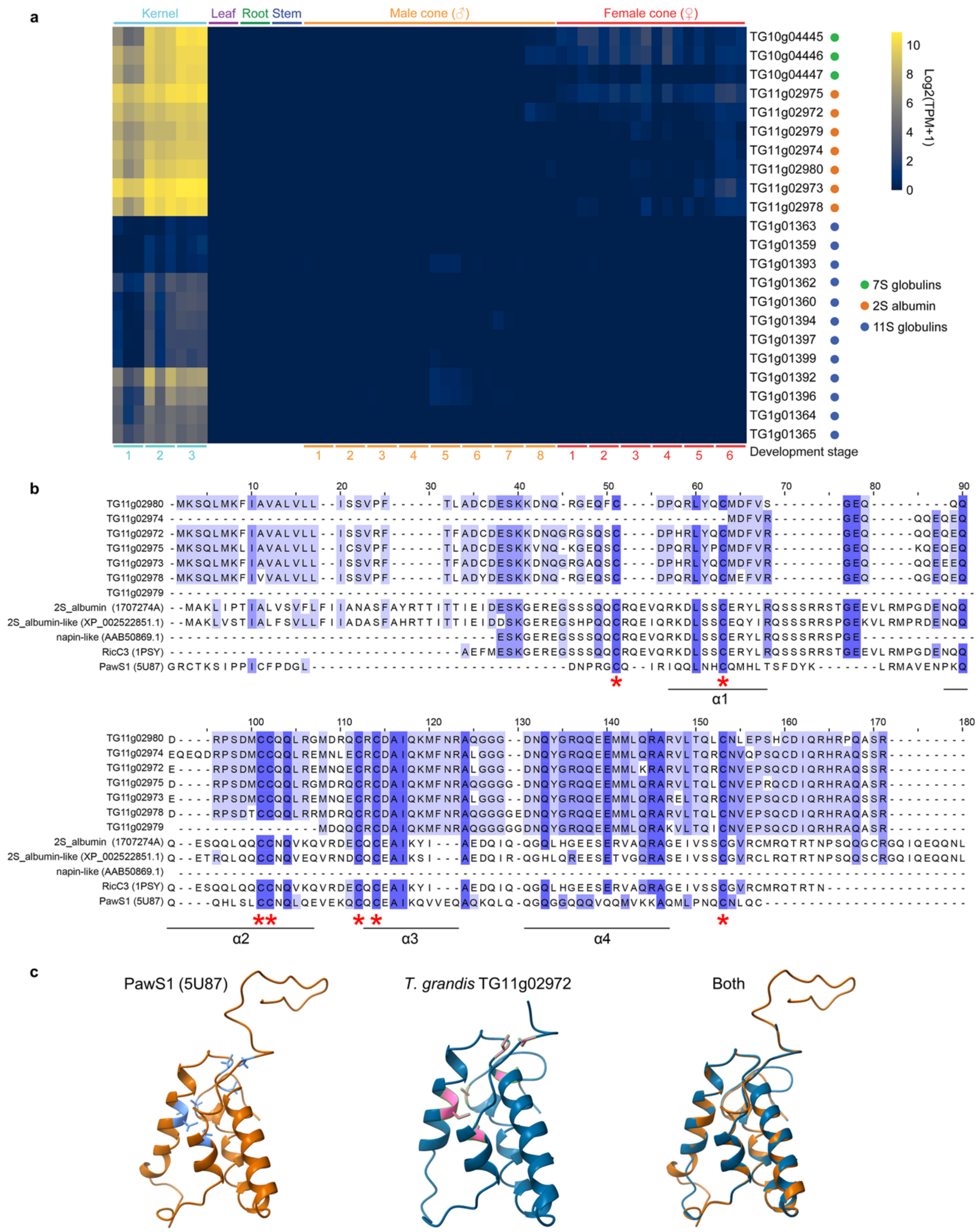


**Supplementary Figure 7. Seed storage proteins in *T. grandis*.** (**a**) Expression of genes encoding seed storage proteins in vegetative and reproductive organs of *T. grandis*. (**b**) Sequence characteristics of 2S albumin family proteins. Red asterisk indicates conserved cysteines. Protein sequences forming the α-helices are underlined. (**c**) Homology modeling of 2S albumin proteins from *T. grandis* and sunflower (PawS1).

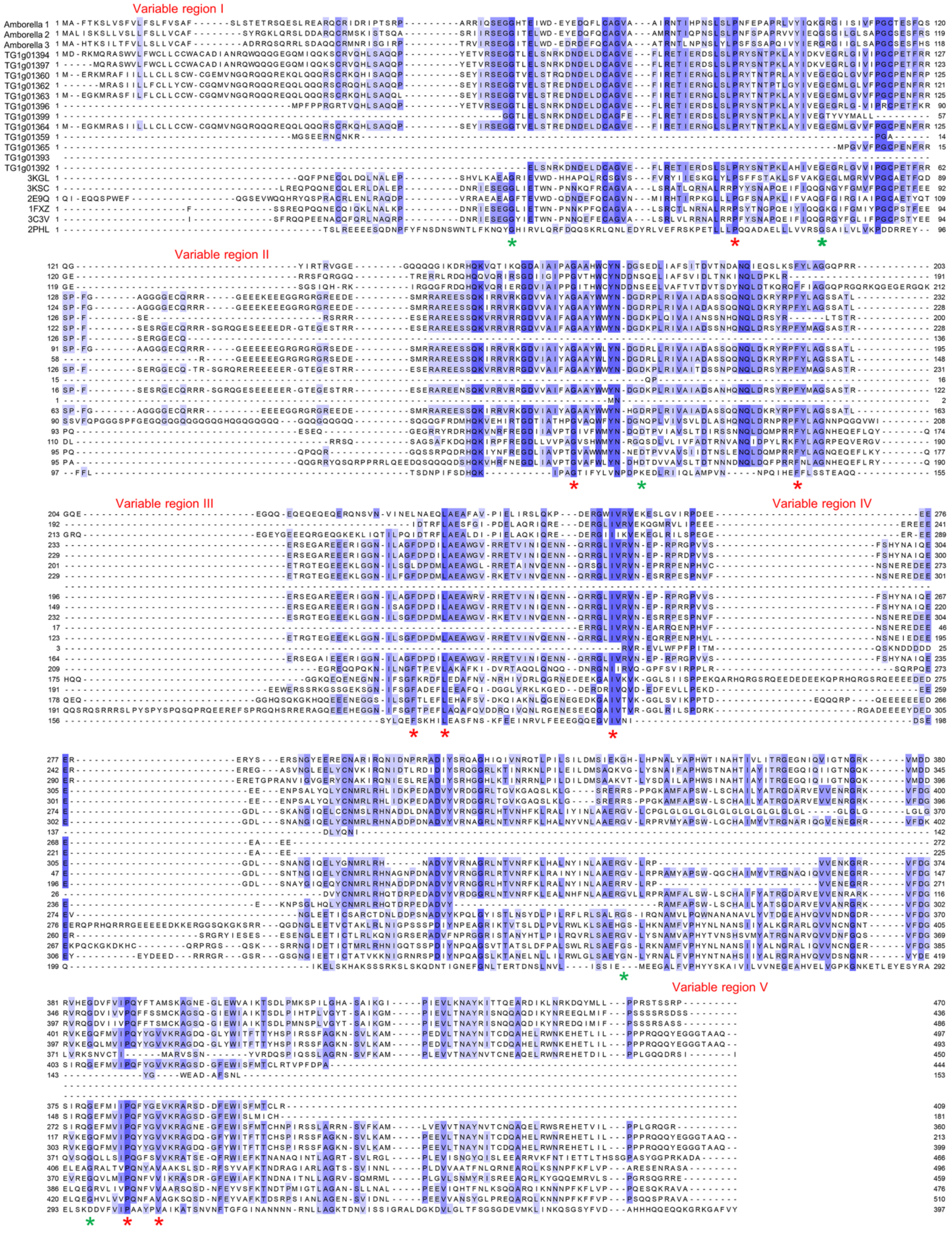


**Supplementary Figure 8. Sequences characteristics of 7S globulins.** Red and green asterisks show functionally important residues that are conserved and variable in *T. grandis*, respectively.

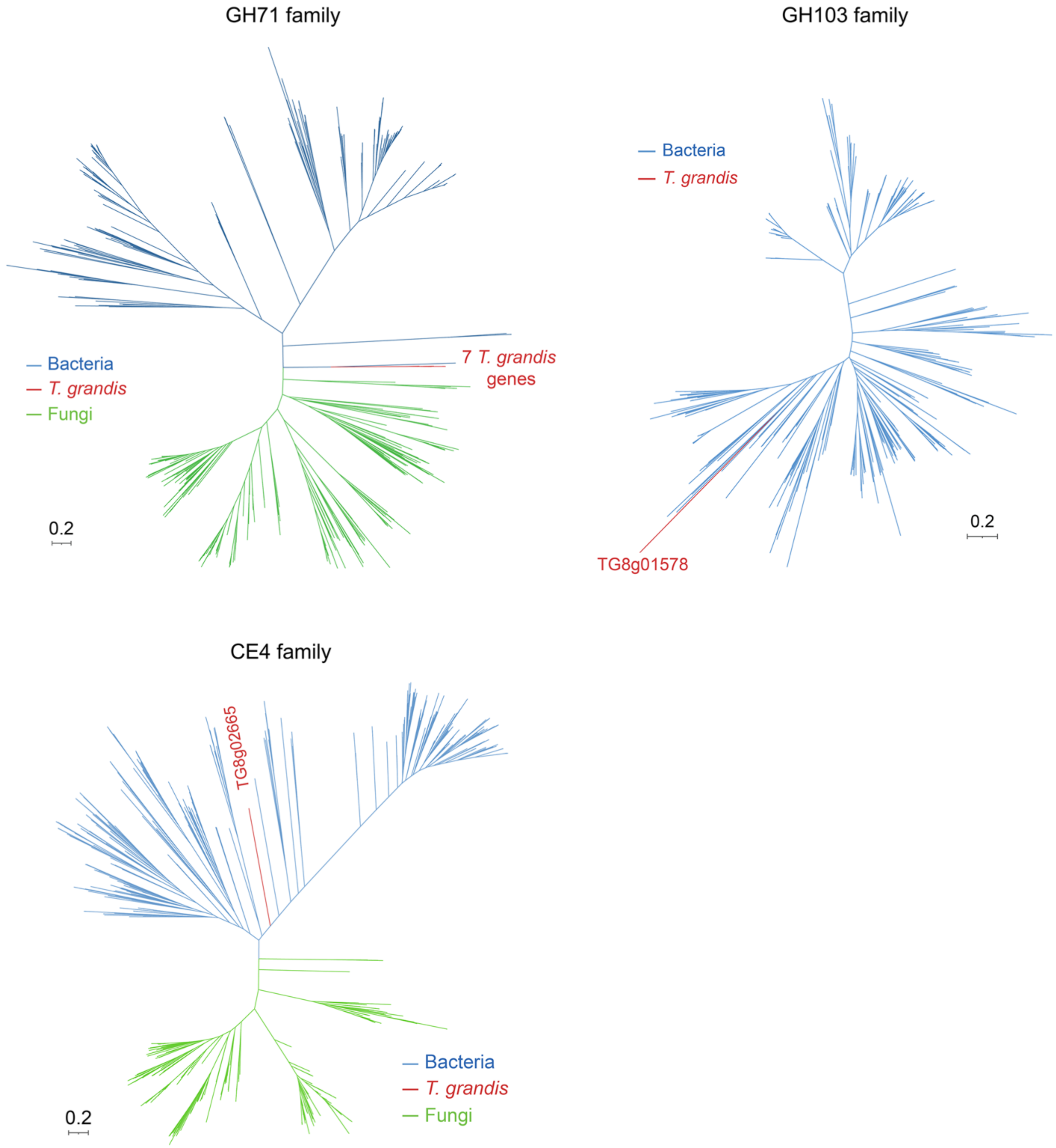


**Supplementary Figure 9. Phylogeny of CAZyme genes.** Protein sequences were aligned using MAFFT with the linsi mode, and the phylogenetic trees were constructed using IQ-Tree with the best-fitting models (Q.pfam+I+G4 for all three families) and 1000 bootstrap replicates.

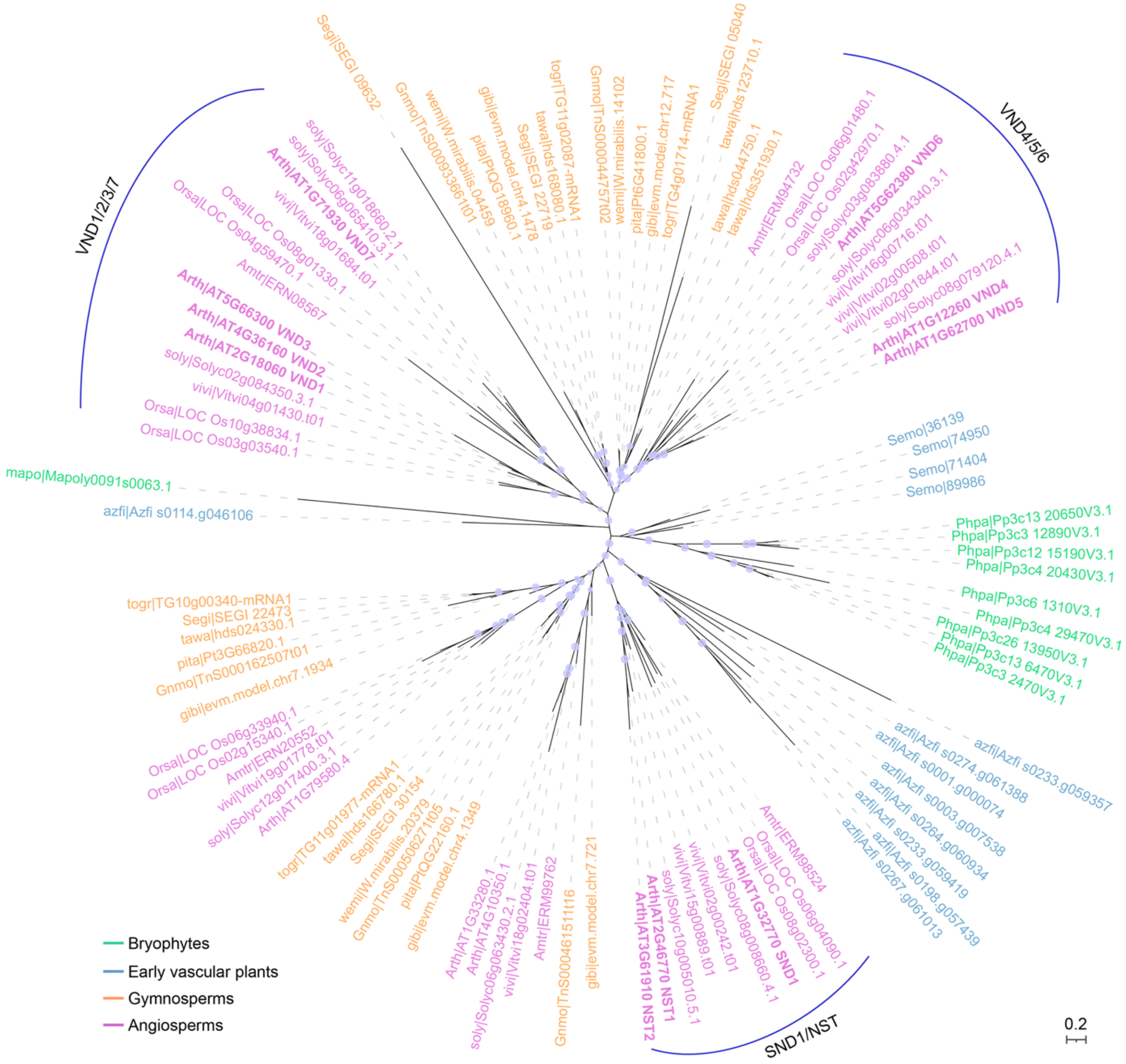


**Supplementary Figure 10. Phylogeny of VND, NST and SND family genes.** Protein sequences were aligned using MAFFT with the linsi mode, and the phylogenetic tree was constructed using IQ-Tree with the best-fitting model (JTT+I+G4) and 1000 bootstrap replicates. Branches with bootstrap support greater than 80% were labeled with light blue pies.

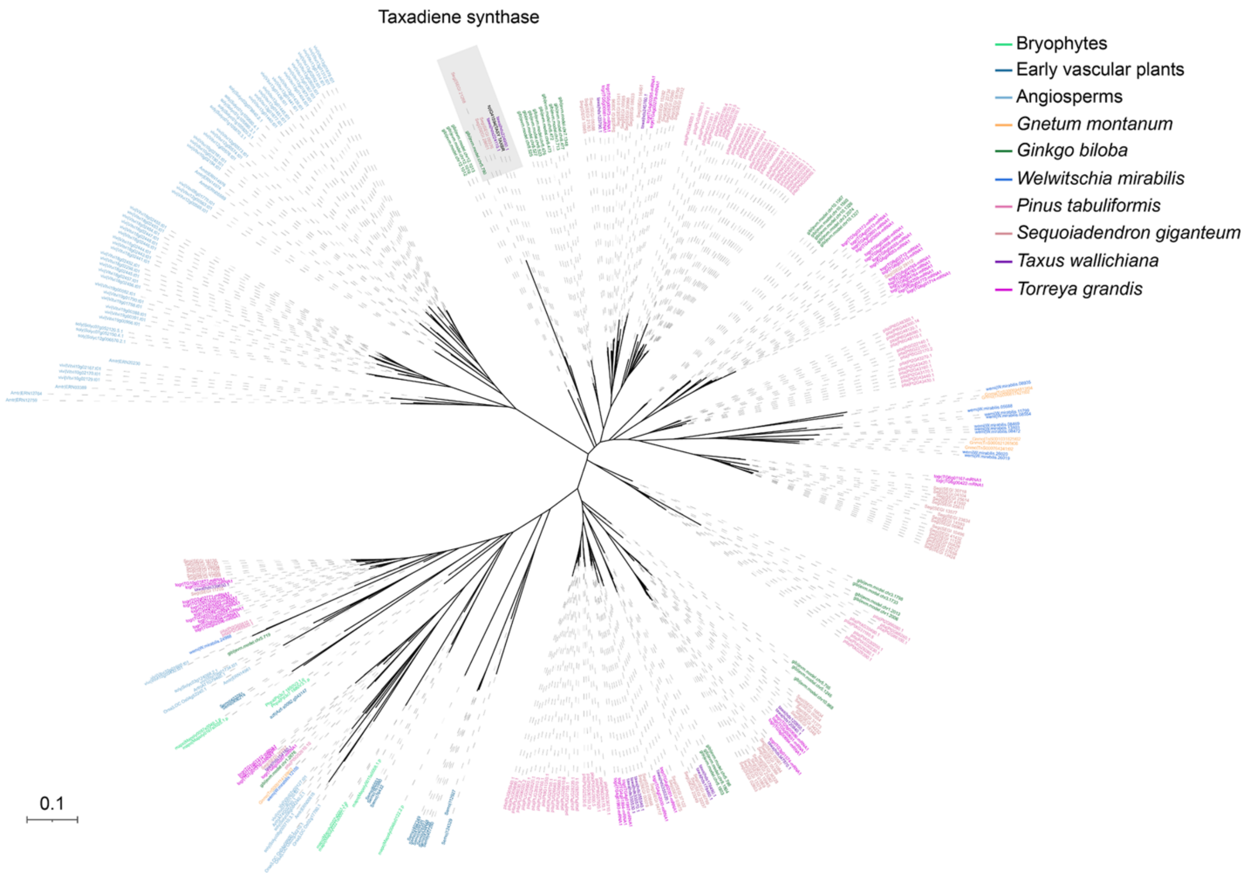


**Supplementary Figure 11. Phylogeny of taxadiene synthase and its close homologues.** Protein sequences were aligned using MAFFT with the linsi mode, and the phylogenetic tree was constructed using IQ-Tree with the best-fitting model (JTT+I+G4) and 1000 bootstrap replicates.

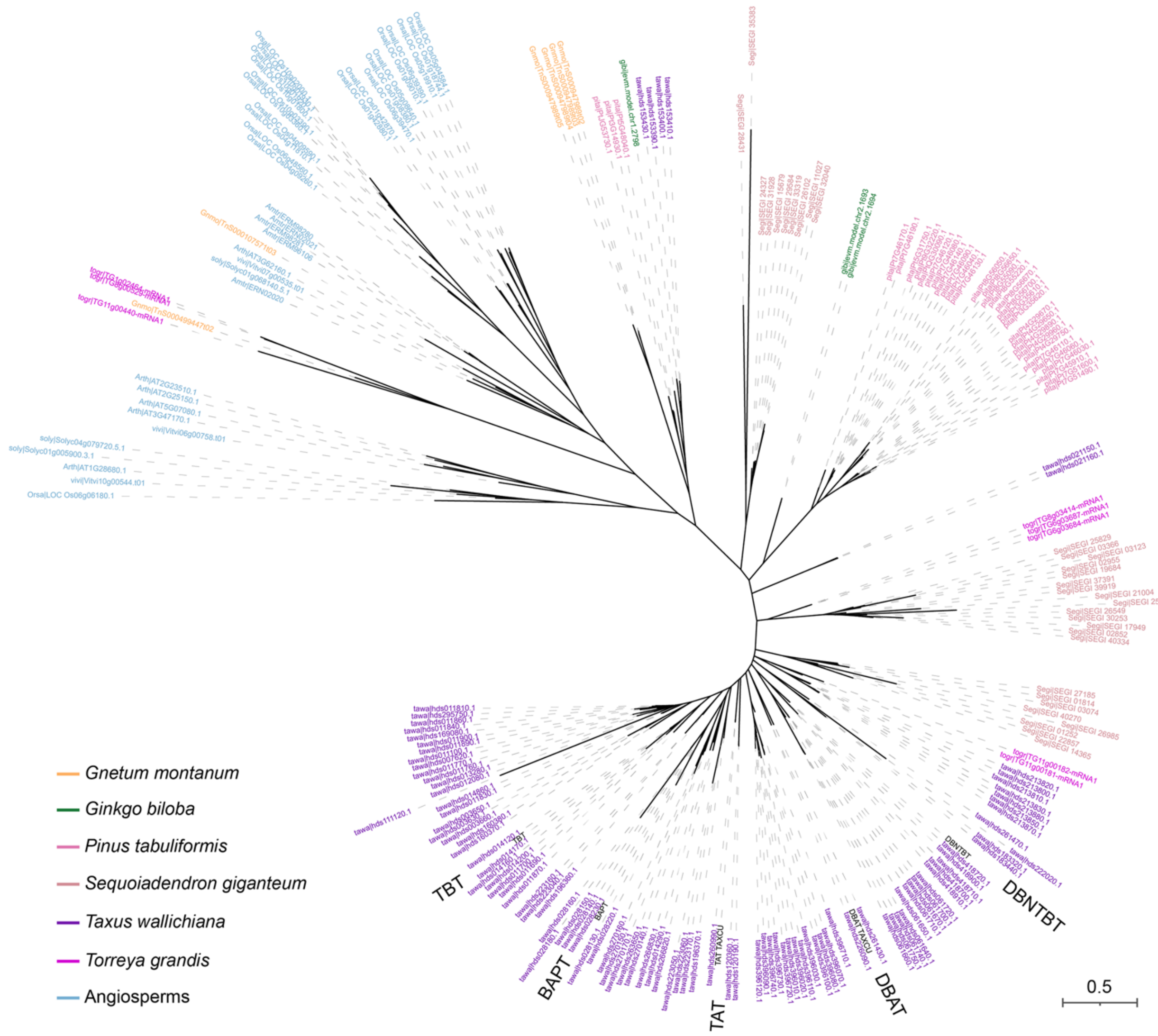


**Supplementary Figure 12. Phylogeny of acetyltransferases.** Protein sequences were aligned using MAFFT program with linsi mode, and the phylogenetic tree was constructed using IQ-Tree with the best-fitting model (JTT+F+G4) and 1000 bootstrap replicates.

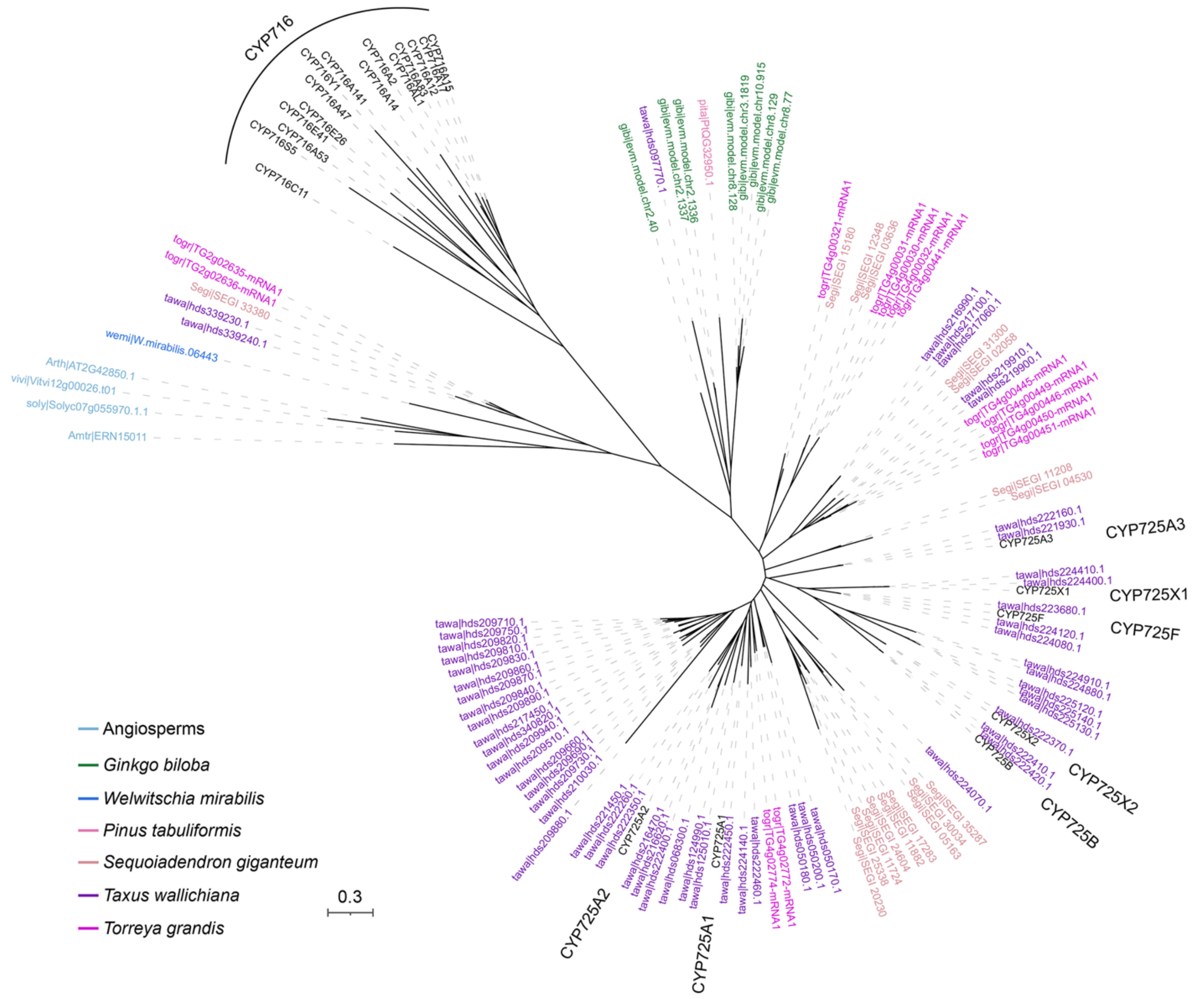


**Supplementary Figure 13. Phylogeny of P450s.** Protein sequences were aligned using MAFFT program with linsi mode, and the phylogenetic tree was constructed using IQ-Tree with the best-fitting model (Q.plant+G4) and 1000 bootstrap replicates.

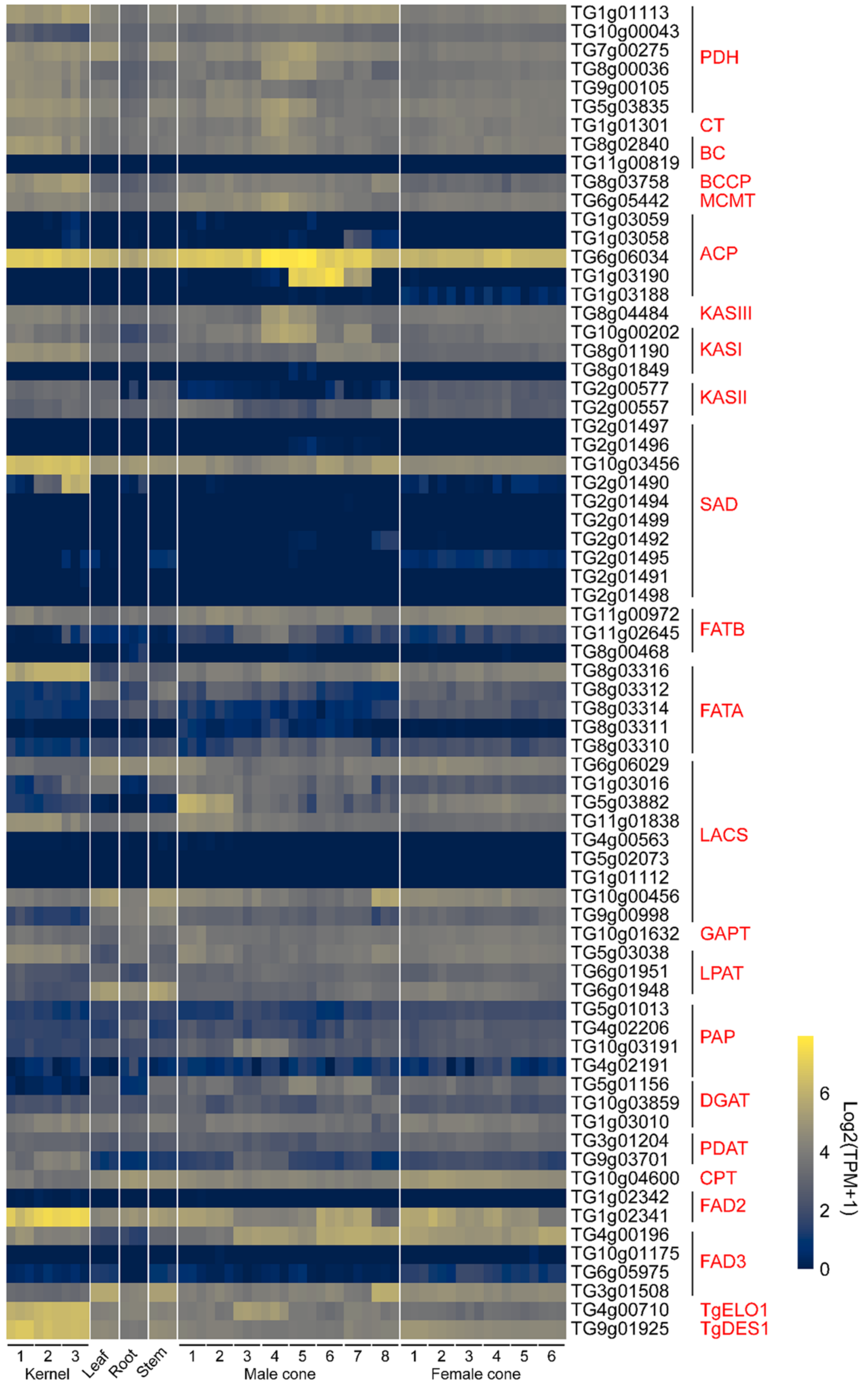


**Supplementary Figure 14. Expression of fatty acid biosynthetic genes in different tissues of *T. grandis*.**

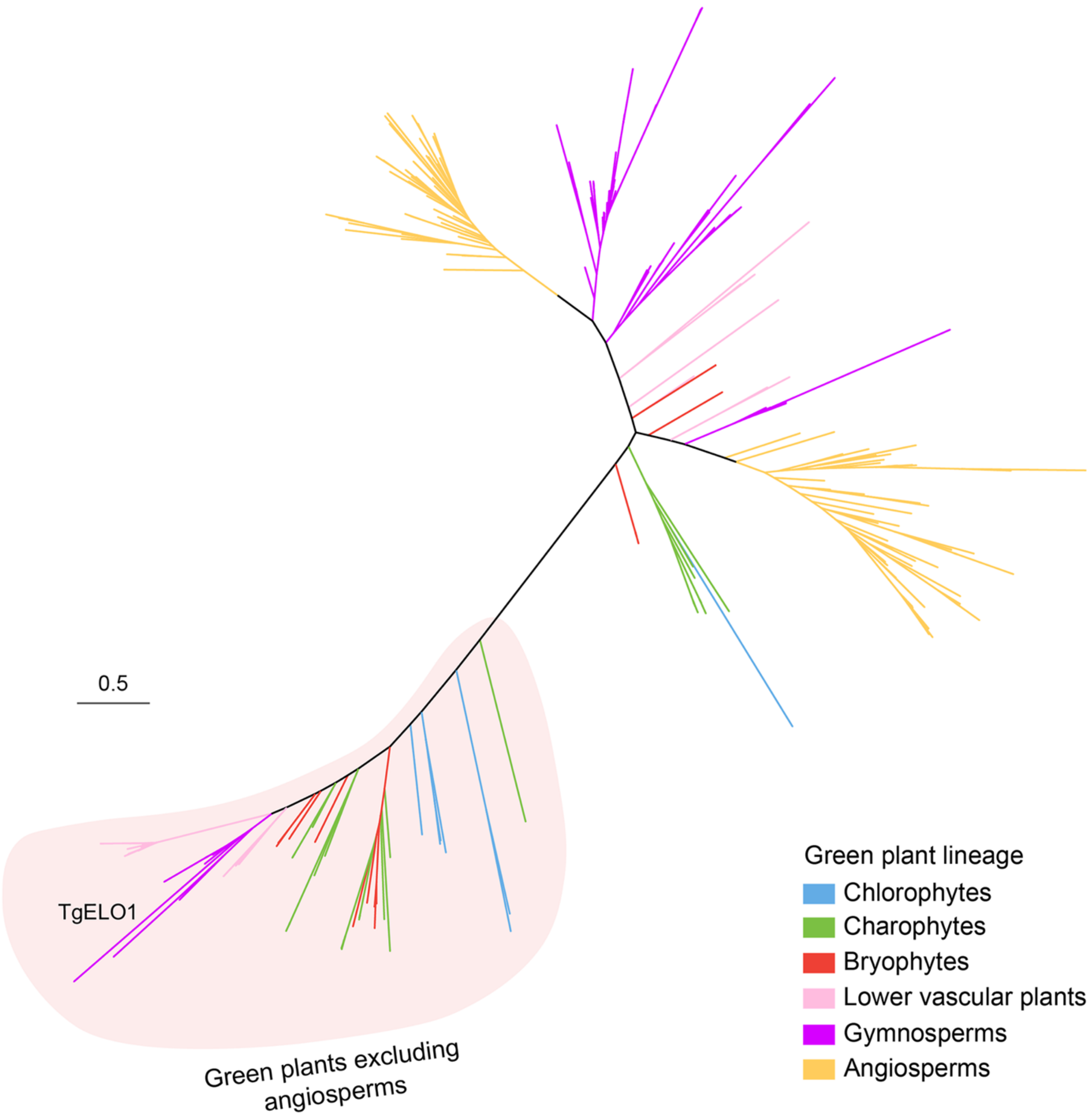


**Supplementary Figure 15. Phylogeny of elongases.** Protein sequences were aligned using MAFFT with linsi mode, and the phylogenetic tree was constructed using IQ-Tree with the best-fitting model (JTT+F+G4) and 1000 bootstrap replicates.

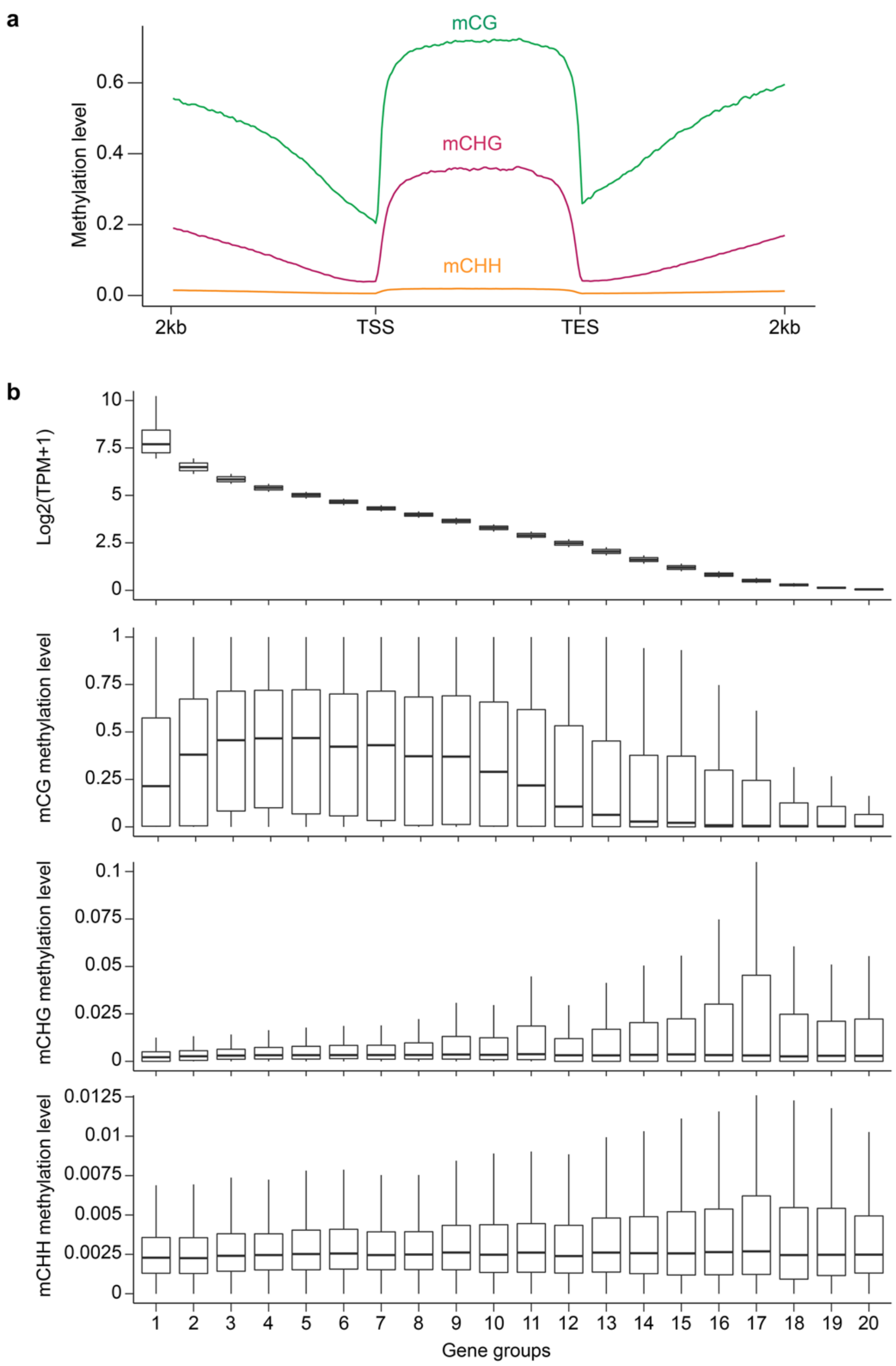


**Supplementary Figure 16. DNA methylation of the seed genome.** (**a**) Methylation levels in gene bodies and the 2-kb flanking regions. TSS, transcriptional start site. TES, transcriptional end site. (**b**) Comparison of gene expression and DNA methylation. Genes were categorized into 20 groups based on the order of their expression levels. For each group, a boxplot was used to show the distribution of gene expression (top) and methylation levels at three different cytosine contexts (bottom).

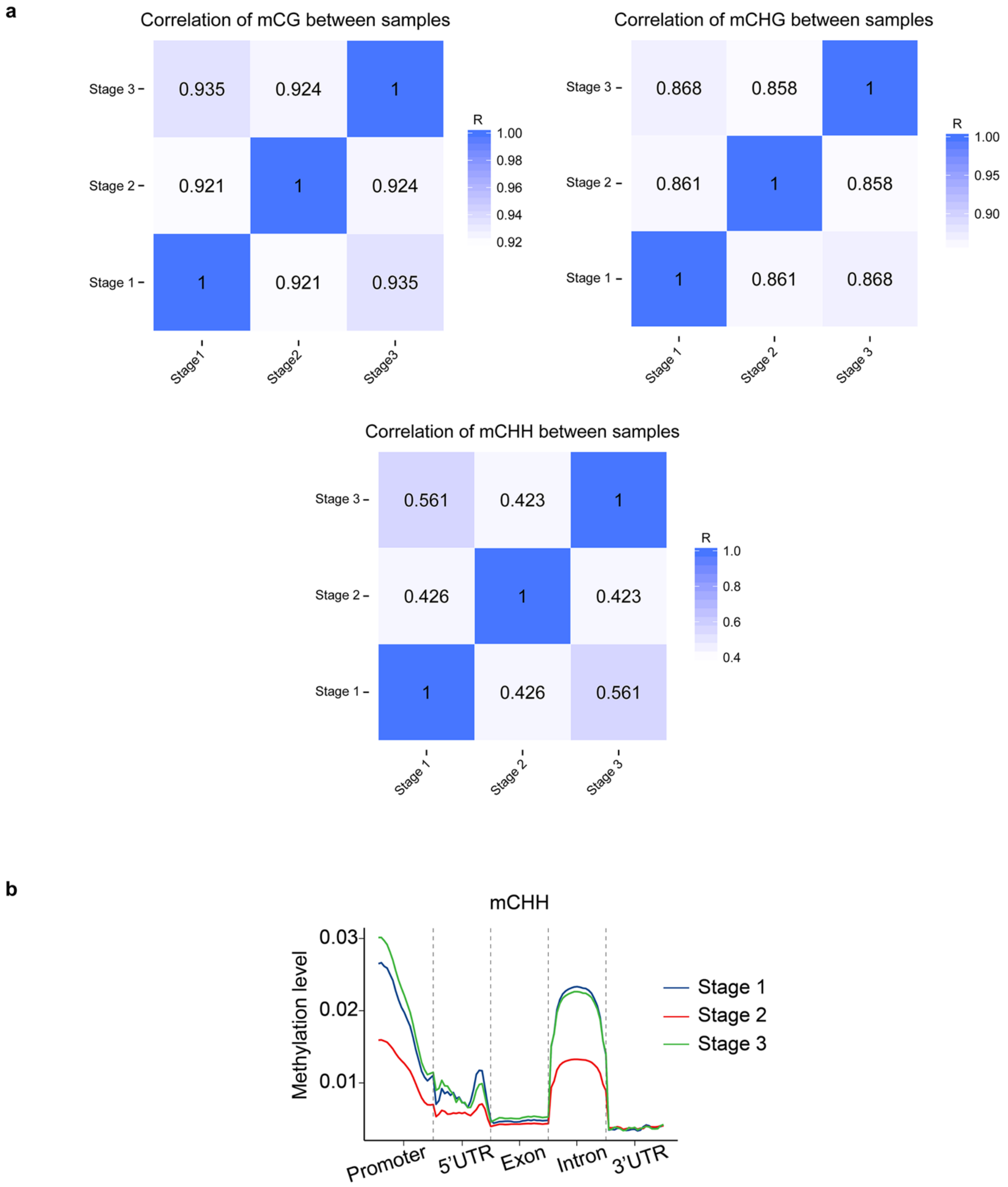


**Supplementary Figure 17. DNA methylation at different stages of seed development.** (**a**) Correlation of methylation levels among different seed development stages. (**b**) mCHH methylation levels of different genomic features in seeds of *T. grandis*

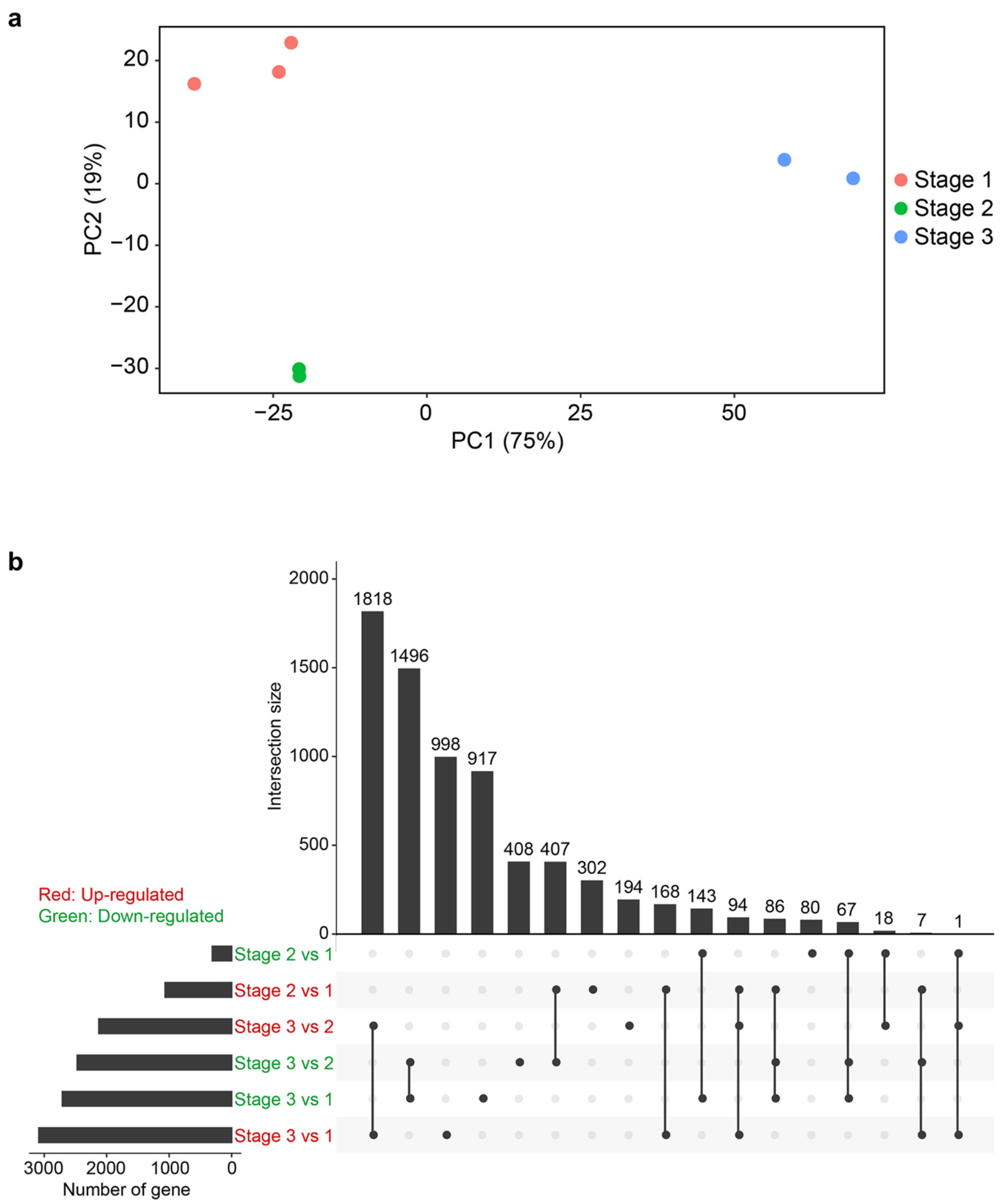


**Supplementary Figure 18. Gene expression profiles at different seed development stages.** (**a**) PCA plot of seed transcriptomes at three development stages. (**b**) Overlaps of differentially expressed genes among three seed development stages.
